# ShearFAST: a user-friendly *in vitro* toolset for high throughput, inexpensive fluid shear stress experiments

**DOI:** 10.1101/2020.01.31.929513

**Authors:** Thomas Brendan Smith, Alessandro Marco De Nunzio, Kamlesh Patel, Haydn Munford, Tabeer Alam, Ohema Powell, Nicola Heneghan, Andrew Ready, Jay Nath, Christian Ludwig

## Abstract

Fluid shear stress is a key modulator of cellular physiology *in vitro* and *in vivo*, but its effects are under-investigated due to requirements for complicated induction methods.

Herein we report the validation of ShearFAST; a smartphone application that measures the rocking profile on a standard laboratory cell rocker and calculates the resulting shear stress arising in tissue culture plates.

ShearFAST measured rocking profiles were validated against a graphical analysis and also against measures reported by an 8-camera motion tracking system.

ShearFAST angle assessments correlated well with both analyses (r ≥0.99, p ≤0.001) with no significant differences in pitch detected across the range of rocking angles tested.

Rocking frequency assessment by ShearFAST also correlated well when compared to the two independent validatory techniques (r ≥0.99, p ≤0.0001), with excellent reproducibility between ShearFAST and video analysis (mean frequency measurement difference of 0.006 ± 0.005Hz) and motion capture analysis (mean frequency measurement difference of 0.008 ± 0.012Hz) These data make the ShearFAST assisted cell rocker model make it an attractive approach for economical, high throughput fluid shear stress experiments.

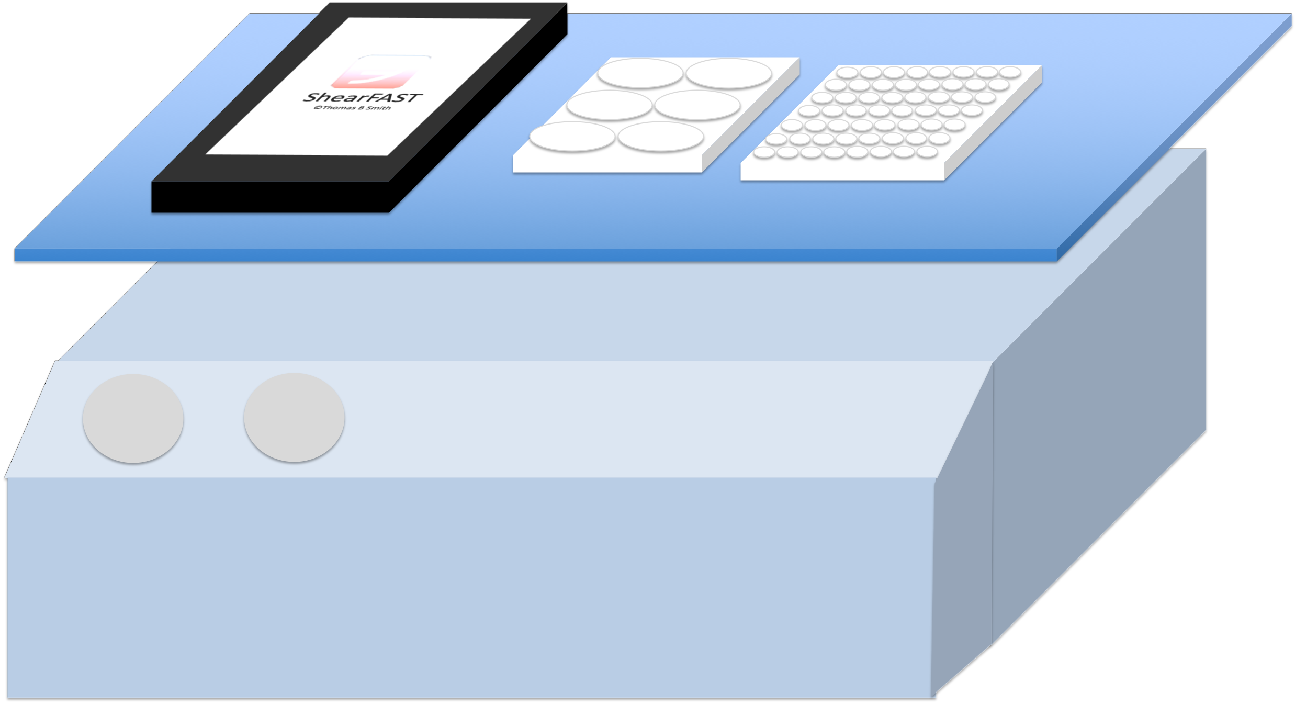

## Introduction

Fluid shear stress (FSS) is described as the deforming force generated against a solid boundary by motile fluid. As an inescapable consequence of fluid flow, FSS is known to modulate cellular physiology in diverse cell types (1–7) and is a key aetiological (8–13), prognostic and therapeutic determinant in multiple disease states (14–18). Despite this, the impact of this key environmental component on cellular physiology remains underrepresented in the majority of cell line based research.

This became apparent in our studies of the impact of machine perfusion techniques on renal tissue destined for transplantation. The presence of fluid flow via low pressure (30 mmHg) pulsatile perfusion is a definitive difference that separates hypothermic machine perfusion (HMP), a mode of *ex-vivo* organ preservation, from more traditional static cold storage techniques. Preservation using HMP results in more favorable post-transplant outcomes when compared to static storage (19–22), and the exertion of fluid flow is a likely mechanism by which HMP promotes such benefit (23–25). However, the optimal HMP environment, including the optimal degree of FSS for each structure within the kidney is yet to be defined by animal models or clinical trials. Finding a solution to this problem may be facilitated through the use of high throughput *in vitro* models.

Cell lines potentiate high throughput screening tools to help direct refinement of organ preservation protocols (26–28), however are generally performed under the absence of fluid flow. This is reflective of standard lab practice; multi-well dishes are an efficient means to subject cultured cells to large numbers of different environments and in general, experiments are performed using assays and equipment compatible with the multi-well plate format.

Although tools exist that allow for simulation of fluid shear stress *in vitro* (29–33), these require bespoke equipment and are limited by their complicated, low throughput, expensive or unscalable nature. These restrictions become particularly pronounced when large scale shear stress experiments are desired.

A cell rocker based method for the delivery of defined degrees of FSS has been described (34) and is utilised in several reports (4,35–39). This approach uses a mathematical model to calculate the resulting fluid shear stress when the rocking parameters (i.e. angle and speed), fluid parameters (i.e. volume and viscosity) and plate dimensions are known.

Many cell rockers possess with a means to adjust rocking angle or speed; however our experience has demonstrated that when even when rocking profile is modifiable, the setting selected may either be incompatible with the model, lack the resolution required or be grossly inaccurate (Figure 6).

Difficulties in delivering the mathematical and analytical accuracy required for the proper execution of cell rocker FSS models may help explain the underutilisation of this otherwise accessible tool in in biomedical research.

Fortunately, the technology to address these problems is currently in place in laboratories throughout the world. Competition between major smartphone manufacturers has led to the ubiquitous presence of handheld devices capable of assessing spatial orientation (40) and performing complex mathematical operations. Since the cell rocker-based approach does not require additional equipment other than the tissue culture dish itself, using smartphones to measure rocking profiles is a simple intuitive step that enables greater experimentation with FSS in cell line studies.

This paper describes the validation of ShearFAST (Shear **<u>F</u>**ormula, **<u>A</u>**ngle, **<u>S</u>**peed **<u>T</u>**oolset), a novel smartphone-based application which enables rapid characterisation of the rocking profile set on a standard laboratory cell rocker, and integrates its findings into the well-established mathematical model of cell rocker based FSS induction.

## Methods

### ShearFAST

ShearFAST is composed of three individual tools; The formula tool, which calculates the characteristic shear stress when experimental parameters are known, the angle tool which measures the maximal rocking angle set on the cell rocker and the speed tool which measures whole cycle rocking frequency.

### Validation of the ShearFAST Formula tool

The formula tool calculates the characteristic fluid shear stress when the volume of fluid, dish diameter, cycle time, fluid volume and viscosity are known. The results of the formula tool (Figure 4) were validated against the example data from the original publication.

### Validation of the ShearFAST angle tool

A graphical analysis method involved capturing side-on photos of the cell rocker, and measurement of the cell rocker platform angle with respect to 0° using ImageJ (41) (Figure 1). A second validation utilised an Infrared Optolectronic 8-camera System (BTS Bioengineering, Milan, Italy) from now on referred to as motion capture analysis (Figure 2).

**Figure 1.**
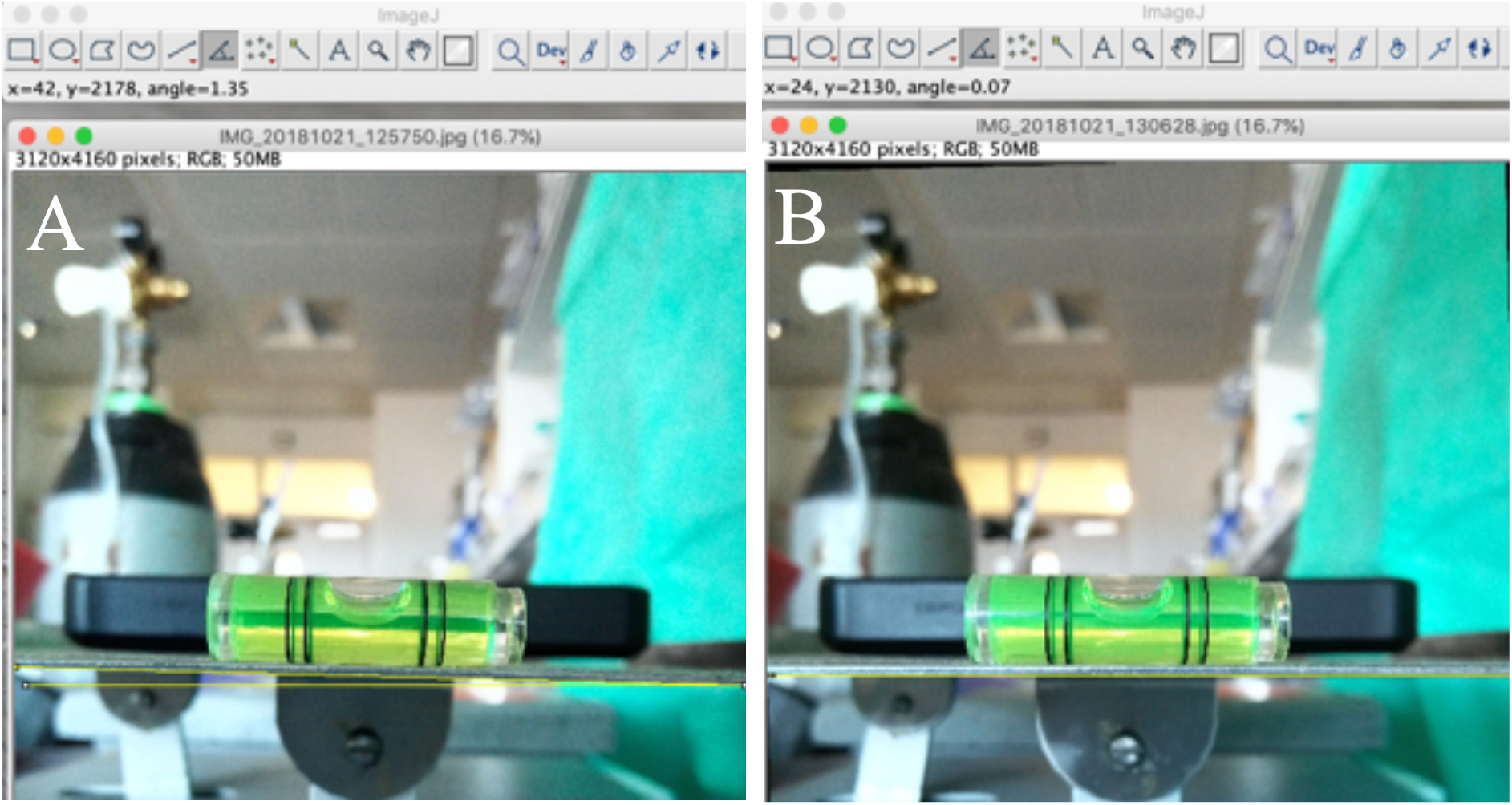
ShearFAST calibration using a spirit level. The leveled platform (A) was used to determine the angle by which the camera capturing the photo was itself offset (B), allowing compensation during ImageJ platform angle analysis.

**Figure 2.**
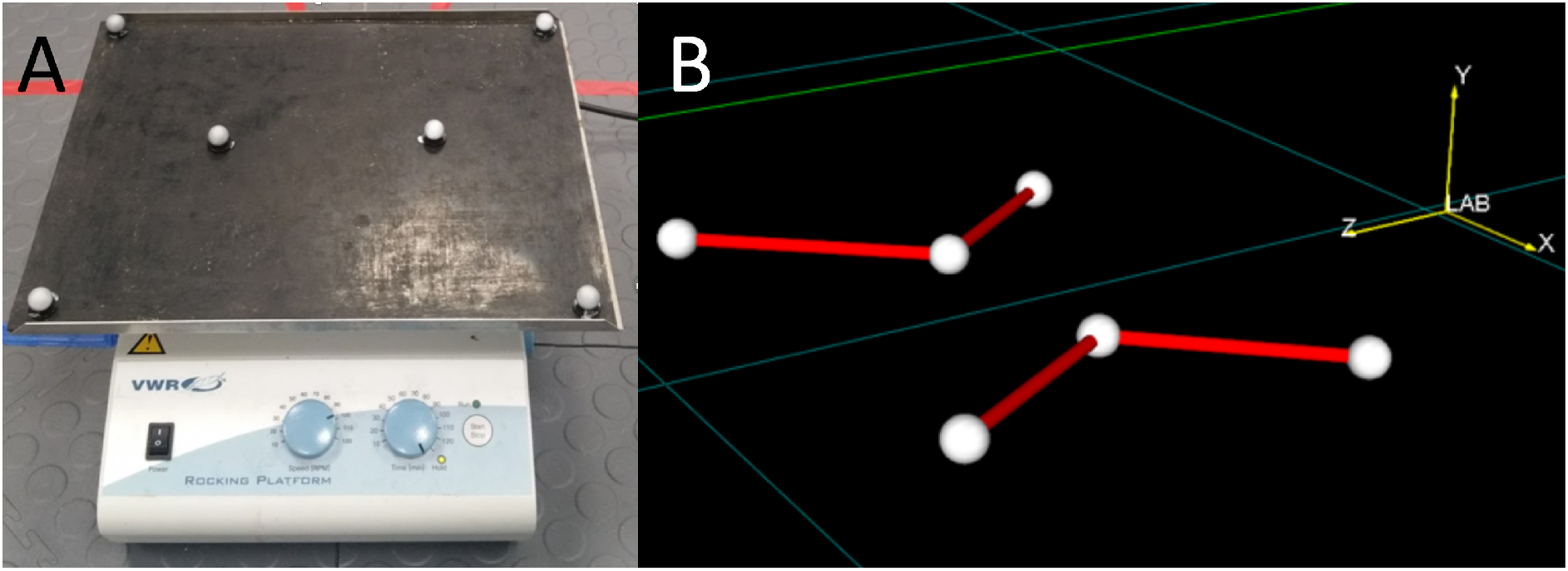
(A) A VWR cell rocker (Cat: 444-0146) which allows manual manipulation of rocking angle and RPM. Six tracking balls were attached to permit detection of rocking angle and cycle frequency. (B) The changes in the geometric position of each tracking ball (i.e. *x,y,z* coordinates) was determined using the 8 camera BTS motion tracking system.

The 2.2 Megapixel infrared cameras resolution (2048×1088 px, BTS DX 6000 model) tracked the 3D motion of retroreflective markers placed on the rocking platform (1.2 cm in diameter) with a precision of 0.1 mm (see https://www.btsbioengineering.com/products/smart-dx-motion-capture/).

### Image J angle analysis

A camera was placed aligned so it could capture a side on view of the cell rocker platform. A smartphone with ShearFAST installed on it was placed on the rocker and the angle tool calibrated to 0° with the aid of a physical spirit level.

An image was taken of the rocker during calibration to allow for later correction of any slight rotation of the camera (Figure1A)

Following calibration, the rocker angle is considered to be 0° as per the spirit level measurements, therefore any apparent rotation of the rocking platform was assumed to be due to a slight camera rotation. The degree required to rotate the image so the platform is perfectly horizontal was determined and applied to the rest of the images after being deemed a suitable approach tested on a second, rotated calibration image (Figure 1B).

After calibration, the cell rocker was set to a series of smartphone detected angles, with screenshots taken of the mean angle detected alongside side view images of the cell rocker. The angles set on the cell rocker images were assessed using the same method, and compared to the measurements obtained using the ShearFAST angle tool.

In a typical cell culture experiment using 35mm plates and 1.5ml media, the authors of the original model recommend avoidance of rocking angle above 10.2° to prevent exposure of the adherent cells to atmosphere. Therefore, 7 angles below this were assessed using the ShearFAST angle tool (Table S1).

### Motion capture analysis

To perform motion capture analysis, the cell rocker was placed at the focal point of an 8-camera motion tracking system. Motion capture retroreflective markers were placed on the rocker (Figure 2A), alongside the smartphone housing the ShearFAST and a physical spirit level. The application was calibrated to 0° as before, then the maximal platform rocking angle adjusted to 12 angles falling between 0-10° using the smartphone application (Table S2). The platform was then set rocking, and mean angle measurements were determined using the SMART ANALYZER software (BTS Bioengineering, version: 1.10.469.0) throughout the rocking period (Figure2B)

### Validation of the Speed tool

#### Video Frequency analysis

The smartphone was placed on cell rocker, which was set to rock at speeds ranging between 30 and 110 rotations per minute (RPM) using dials on the rocker. Using ShearFAST, the acquired waveform was compared to modelled waveforms of user defined frequency until the acquired data was overlaid, granting rapid determination of rocking frequency.

A video was taken of each rocking experiment during ShearFAST data acquisition. The videos were observed alongside a stopwatch with millisecond resolution. A screencast of both video and stopwatch was recorded, and the timepoints at which the cell rocker reached its maximal rocking angle was determined over the course of the video (Figure 3).

**Figure 3.**
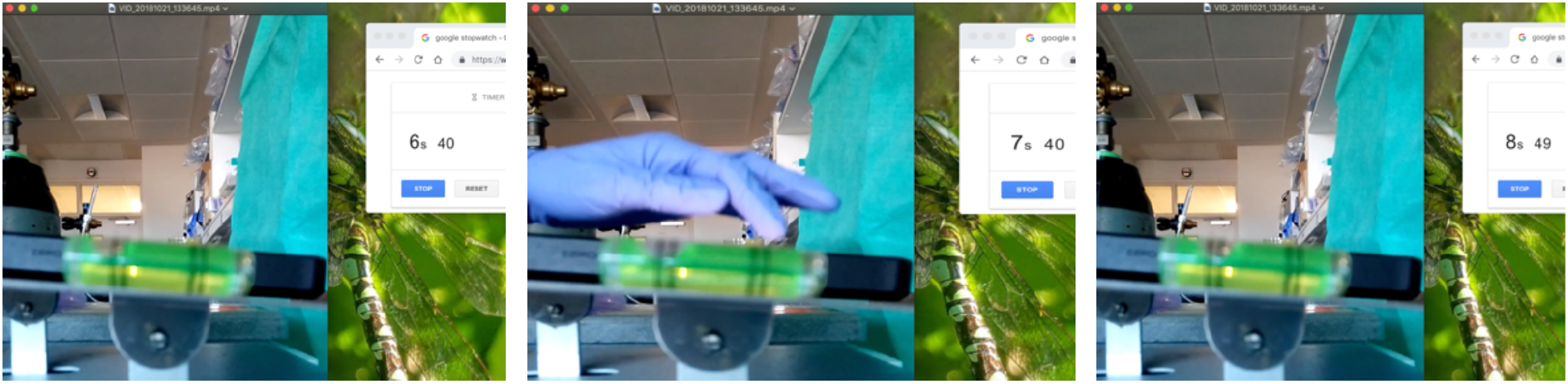
Measurement of cycle time using video analysis. The images show consecutive timepoints at which the rocking platform reached its maximum rocking angle, which were recorded.

At least three timepoints were collected for each rocking speed, subtraction of each timepoint from its preceding timepoint granted the assessment of the mean time taken to complete one full rocking cycle. These were averaged (*T)*, permitting calculation of cycle frequency (*Hz*) using Equation 1. The data is reported in Table 1, and correlations between the measurement methods reported in Figure 7.

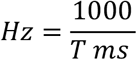

Equation 1: conversion of mean cycle time to rocking frequency simulations and frequency measured during video analysis

**Table 1.**
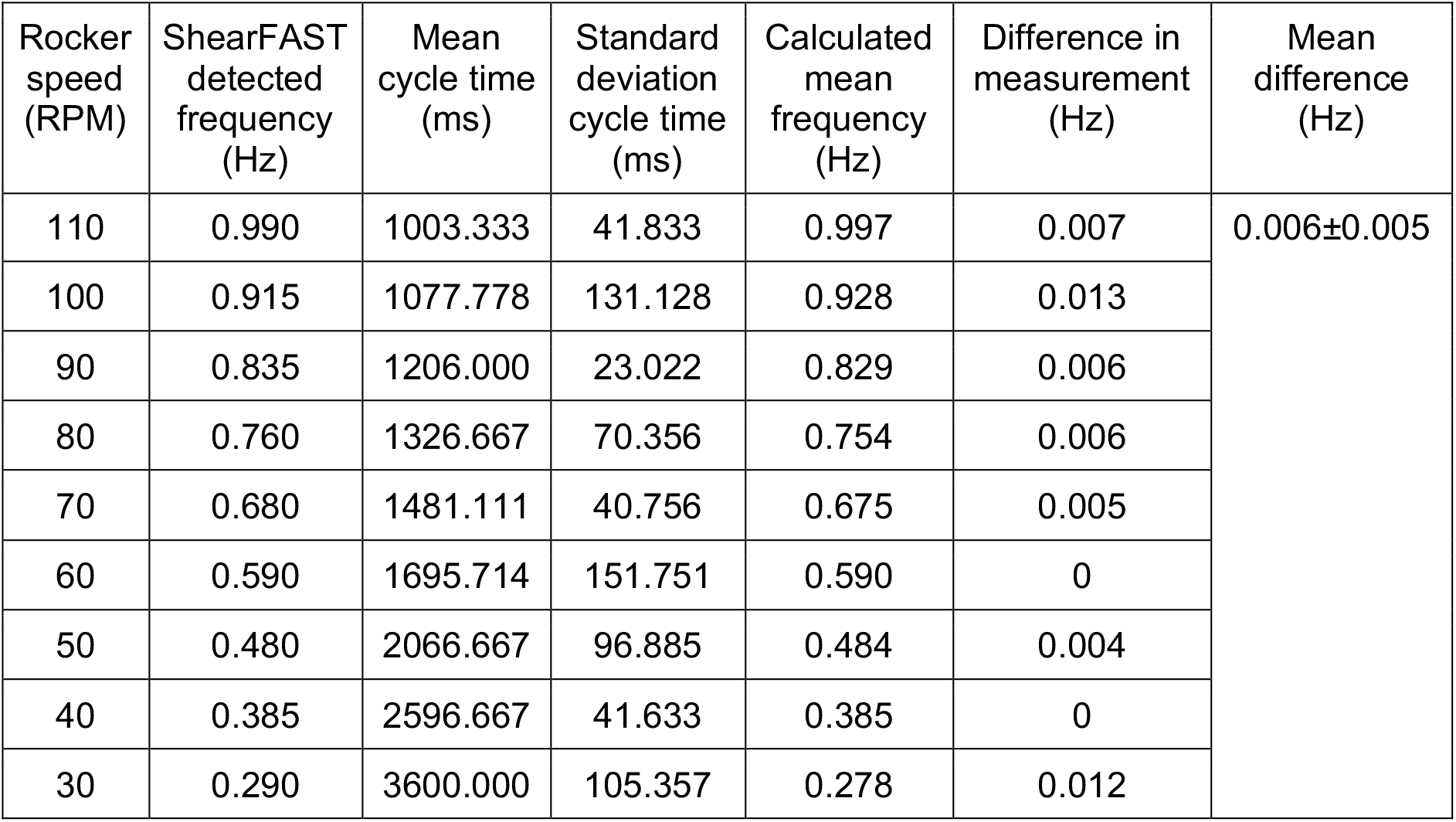
The rocking frequency detected by ShearFAST at different rocking speeds.

| Rocker speed (RPM) | ShearFAST detected frequency (Hz) | Mean cycle time (ms) | Standard deviation cycle time (ms) | Calculated mean frequency (Hz) | Difference in measurement (Hz) | Mean difference (Hz) |
| --- | --- | --- | --- | --- | --- | --- |
| 110 | 0.990 | 1003.333 | 41.833 | 0.997 | 0.007 | 0.006±0.005 |
| 100 | 0.915 | 1077.778 | 131.128 | 0.928 | 0.013 |  |
| 90 | 0.835 | 1206.000 | 23.022 | 0.829 | 0.006 |  |
| 80 | 0.760 | 1326.667 | 70.356 | 0.754 | 0.006 |  |
| 70 | 0.680 | 1481.111 | 40.756 | 0.675 | 0.005 |  |
| 60 | 0.590 | 1695.714 | 151.751 | 0.590 | 0 |  |
| 50 | 0.480 | 2066.667 | 96.885 | 0.484 | 0.004 |  |
| 40 | 0.385 | 2596.667 | 41.633 | 0.385 | 0 |  |
| 30 | 0.290 | 3600.000 | 105.357 | 0.278 | 0.012 |  |

#### Motion capture Frequency analysis

Motion capture frequency analysis granted an assessment of the accuracy and reproducibility of the smartphone frequency measurements. Using the ShearFAST, the rocking frequency was set to 1Hz for three measures, and 0.65Hz for four other measures and compared to the motion capture results (Table 2).

**Table 2.** Reproducible frequency assessment by ShearFAST and motion capture analysis

| ShearFAST determined frequency (Hz) | Motion capture frequency (Hz) | Difference (Hz) | Mean Difference (Hz) |
| --- | --- | --- | --- |
| 1.0 | 1.035 | 0.035 | $0.007 \pm 0.012$ |
| 1.0 | 0.995 | 0.005 |  |
| 1.0 | 0.995 | 0.005 |  |
| 1.0 | 1.0 | 0.0 |  |
| 0.65 | 0.645 | 0.005 |  |
| 0.65 | 0.655 | 0.005 |  |
| 0.65 | 0.65 | 0.0 |  |
| 0.65 | 0.65 | 0.0 |  |

#### Statistical analysis

Correlations between ShearFAST measurements of rocking angle or frequency and those detected by the other analysis were identified by the Spearman R test. Linear regression was used to determine if the slope generated by ShearFAST measurements of rocking pitch or frequency differed from those detected by the other analysis.

## Results

### Validation of the Formula tool

The formula tool calculates the characteristic fluid shear stress reported in when the volume of fluid, dish diameter, cycle speed, fluid volume and viscosity are known (Figure 4). The FSS calculated by the ShearFAST formula tool was validated against the example data from the original publication’s supplementary data (34)

**Figure 4.**
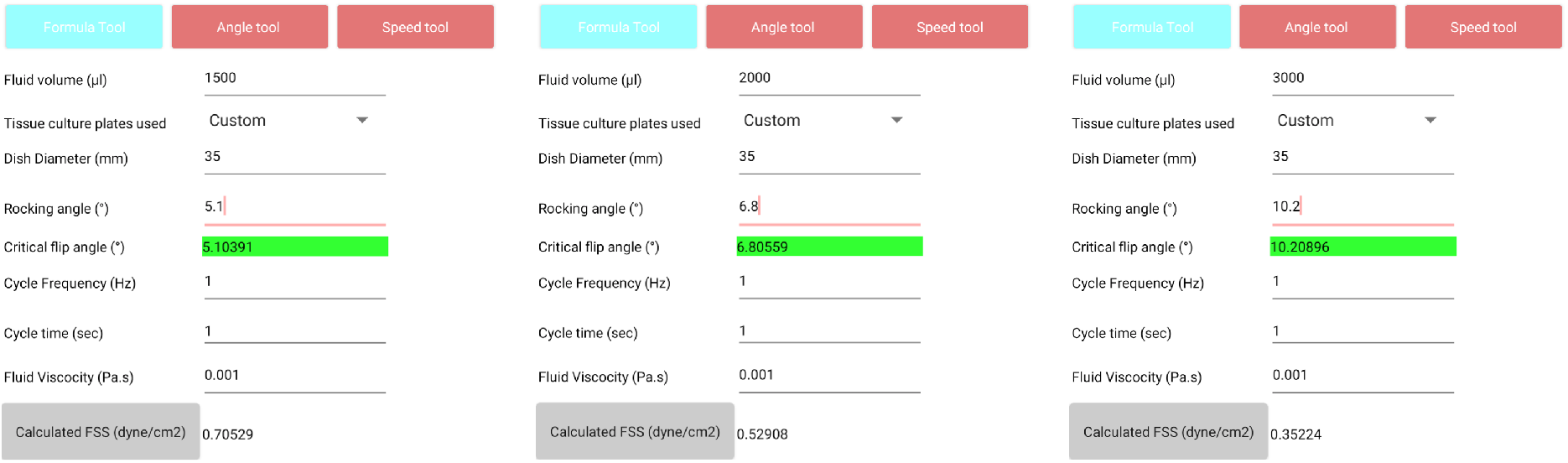
ShearFAST formula tool reproduces the example data reported in the supplementary data of the original publication original publication

### Validation of the Angle tool

#### ImageJ angle analysis

ImageJ angle analysis demonstrated a close association with the smartphone application measurements (r=1.0, p≤0.0004) (5A). The smartphone measurements were around 0.25° higher than those of the graphical analysis (Table S1), but no difference in slope was found between angle measurement methods (p>0.84).

#### Motion capture angle analysis

On average, ShearFAST angle measures and those attained by motion capture analysis varied by 0.54 ± 0.16° however this difference was not found to be significant (p>0.32) (Table S2). Angle measures between the two angle assessment methods were found to be correlated (R=0.99, p ≤0.0001) (Figure 5B).

**Figure 5.**
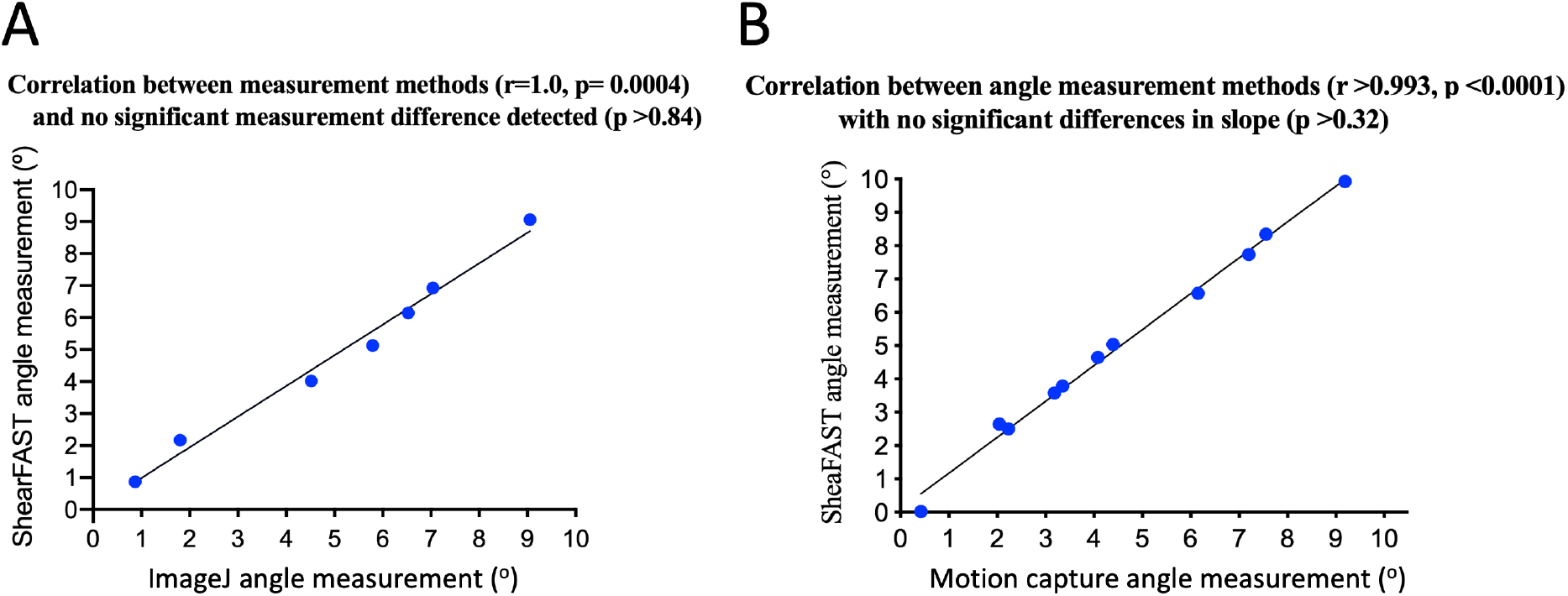
Correlation between pitch measurements detected using ShearFAST and Pitch measurements detected using ImageJ(A) and motion capture analysis (B)

#### Video frequency analysis

Video analysis of rocker cycle frequency corroborates the frequency assessment by the smartphone application (Table 1) and (Figure 6), with strong association between measurement techniques (r≥0.99, p≤0.0001) and no significant difference detected between the two measurement techniques (p=0.9219) (Figure 7).

**Figure 6.**
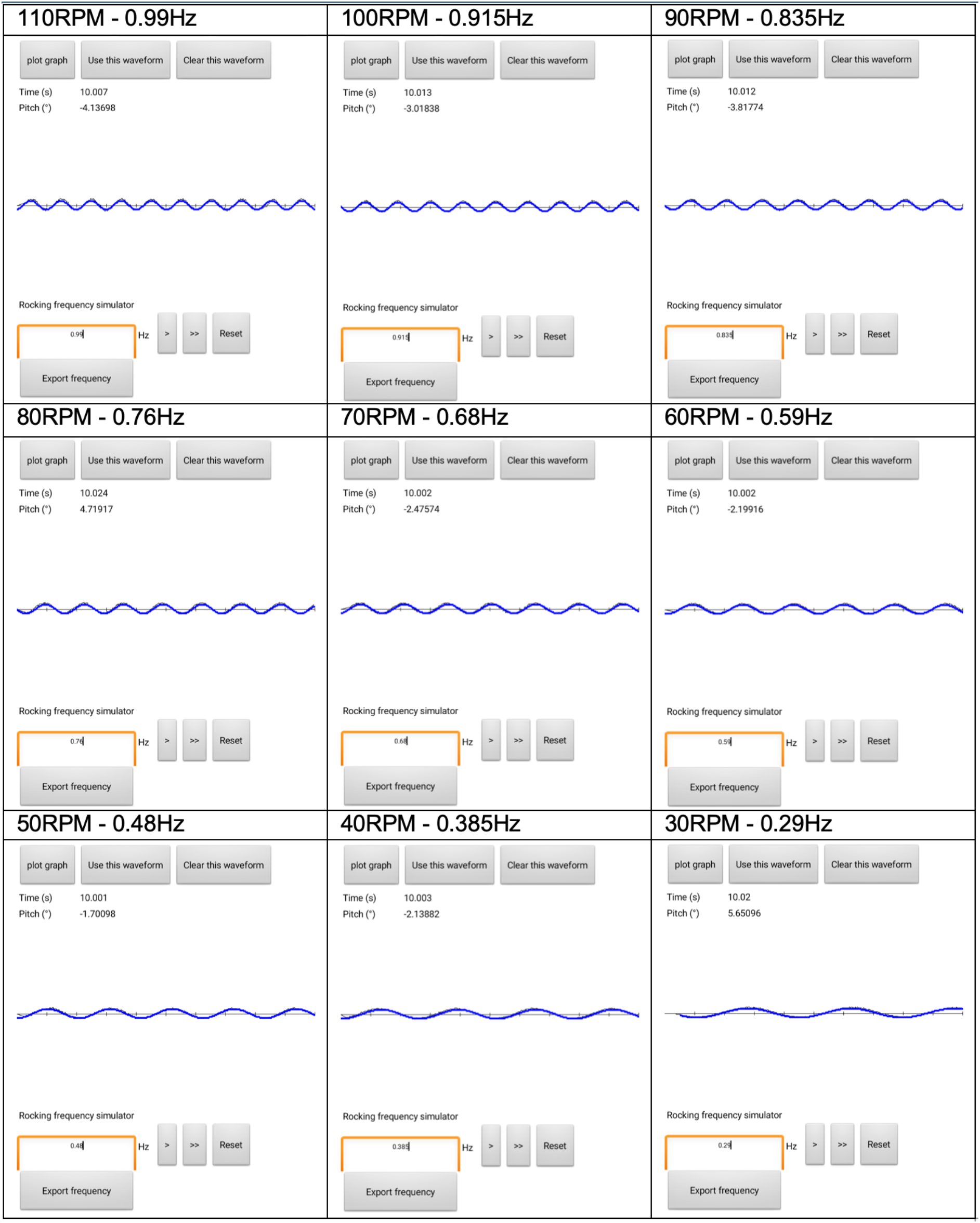
ShearFAST measured rocking frequencies at rocker defined rocking speeds (RPM). Illustrating the value of the ShearFAST approach; when manually selecting the rocking speed 60RPM (i.e. 1Hz) the resulting profile was visibly slower. ShearFAST determined a rocking speed of 0.59Hz which was corroborated by later video analysis (Table 1).

**Figure 7.**
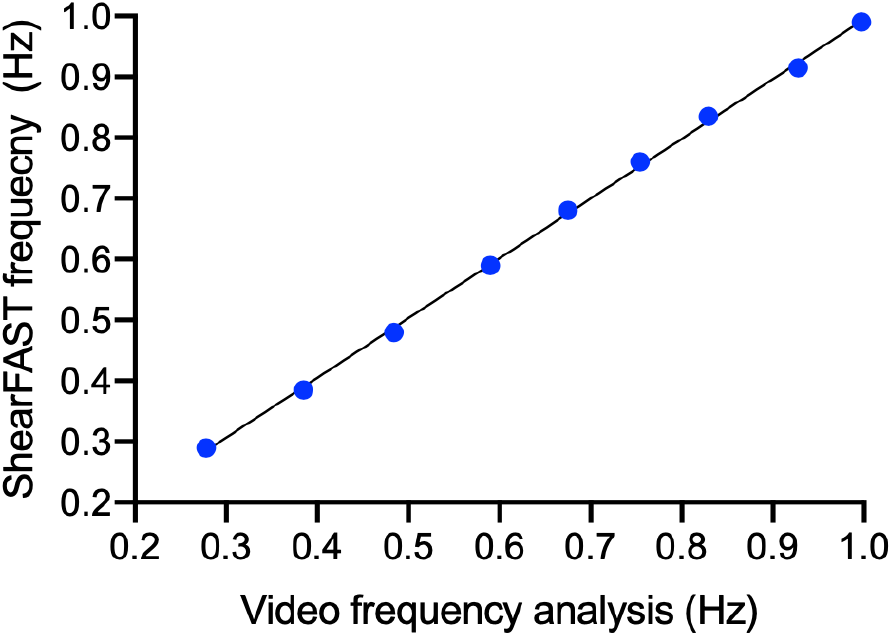
Correlation between ShearFAST frequency measurement and video frequency measurement.

#### Motion capture frequency analysis

This analysis revealed a small average difference in frequency assessment of 0.007 ± 0.012Hz between measures (Table 2).

## Discussion

Within cell line research there is a drive to more accurately simulate physiological environments through the utilization of parallel plate flow chambers, microfluidics and organ-on-a-chip devices. These allow continuous instigation of fluid flow, however arguably diminish the versatility of the cell line approach.

The fixed growth area available to adherent cells cultured in such devices alongside the requirements for sperate perfusion circuits restricts both the type and number of the downstream analysis that can be performed.

The cell rocker model was proposed in 2010 (34), and has been utilised to demonstrate important effects of FSS in diverse cell lineages and areas of medical research; (4,35–39). However, the complexity of the model, including its requirements for mathematical and technical accuracy may be a factor limiting its correct implementation.

ShearFAST provides a user-friendly experience with which to perform high throughput and scalable cell rocker induced cell line experiments in conventional laboratory cultureware. We demonstrate that the speed tool interface is an effective, intuitive and very efficient means for the determination of whole cycle rocking frequency. ShearFAST measures of rocking frequency correlated strongly with those derived from the real time video analysis, with very small, insignificant and ineffectual differences observed when compared to the video and motion capture analysis (i.e. 0.006Hz and 0.007Hz respectively).

We also found the angle tool measures platform pitch with acceptable degrees of accuracy. No differences in slope were detected by between ShearFAST measured angles and motion capture analysis or graphical analysis.

Consider a hypothetical experiment in which 1.5ml of cell culture media is added to a 35mm dish and rocked at 1Hz. On average, the mean imprecision between the ShearFAST angle tool and the techniques used to validate it were found to be 0.54° for the motion capture analysis and 0.29° for the ImageJ analysis. Assuming perfect accuracy by the validatory methods and an angle overestimation by ShearFAST, these mean degrees of imprecision would result in a characteristic shear stress generation of 0.63 dyne/cm^2^ and 0.66 dyne/cm^2^ respectively, rather than the mathematically calculated 0.7 dyne/cm^2^ resulting from the conditions in the described hypothetical experiment.

Since, as modelled in the original rocker publication, FSS induced during seesaw rocking varies with both cycle time and dish location, these insignificant differences detected in pitch measurements are offset by the potential of ShearFAST to foster simple, reproduceable and high throughput FSS experiments.

The ShearFAST FSS induction method is compatible with high throughput apparatus to control additional extracellular environments, such as oxygen availability using hypoxia chambers. We performed the example screen under fluid anoxia using a nitrogen purged hypoxia chamber, thus we can be confident the protective effect of FSS observed did not stem from improved oxygen delivery to the submerged monolayers.

Cultured proximal tubule cells have been shown to respond dynamically to fluid shear stress, changing both their phenotype (44), reabsorbative activities (1) as well as the 3D actin structure of their cytoskeleton (45).

We found the mean differences in cellular viability caused by generation of fluid flux to be small (i.e. instigation of 1 dyne/cm^2^ resulted in a 7% improvement in viability when compared to cells stored under fluid stasis).

This emphasizes the value of the ShearFAST assisted cell rocker model. Unlike alternative methods of FSS induction, it is simple and economical to perform large numbers of technical or biological replicates using the multiwell plate format. Through this advantage, the effects of biological variation between experiments may be silenced, leading to enhanced detection of small changes conferred by fluid movement in the extracellular environment.

Additionally, ShearFAST permits the generation of different degrees of FSS in the same experiment. The volume of fluid used is a variable governing the FSS generated under the same rocking profile. When paired with static parallel plates containing the same volumes of fluid, this phenomenon may be manipulated within ShearFAST to perform a ‘dose response’ of FSS induction.

## Conclusions

ShearFAST is a useful tool which facilitates the rapid execution of fluid shear stress experiments using a cell rocker. The mobile format of the software permits instant user acquisition through established smartphone application providers, while the user interface provides an intuitive means with which to accurately measure the rocking profile on a standard laboratory cell rocker. The costless execution of ShearFAST assisted cell are particularly useful for proof of concept studies, and is applicable for research involving adherent cell lines.

## Acknowledgements

The authors would like to thank Kalliopy Tabry for her invaluable contributions to this work. This work was supported by a generous grant from University Hospital Birmingham Charities.

